# A Viral Mutation Profiling and Discovery Strategy for Sensitive Multiplex Detection of Viruses and Variants in Saliva by Proteomics

**DOI:** 10.64898/2026.05.30.729008

**Authors:** Yanjia Zhang, Michael R. Shortreed, Aaron T. Timperman

## Abstract

**Background:** Detecting multiple viruses in biofluids such as saliva remains challenging because of rapid mutation and low abundance presenting in the complex matrix. Proteomics analysis provides a complementary method by directly measuring viral peptides and resolving protein-level sequence variation in rapidly evolving viruses such as influenza and SARS-CoV-2. However, the ongoing viral evolution challenges search sensitivity by introducing numerous mutations. Here, we develop a saliva viral proteomics strategy that integrates mutation profiling of clinically relevant viral sequences with single amino acid substitution mutation discovery to enable sensitive detection of both known and emerging variants.

**Results:** *In silico* analysis of SARS-CoV-2 and influenza viral sequences reveals bimodal prevalence distributions of mutated peptides. For mutation profiling of clinically relevant viral sequences, we prioritize high-prevalence peptides, which presumably have higher clinical significance, within a 3-month timespan. Applying a 10% prevalence threshold reduces database size by more than 82.1% for SARS-CoV-2 and 94.5% for influenza. Optimized mutated peptide databases cause <0.8% sensitivity loss compared with searches using viral reference sequences alone, while still covering 97.8% of SARS-CoV-2 and 98.4% of influenza viral populations in the subsequent month. G-PTM-D-based mutation discovery further mitigates the loss of database coverage and enables detection of new variants. In peptide-spiked saliva samples, G-PTM-D detected single-amino-acid variant peptides with femtogram-level sensitivity, outperforming conventional PTM searches and *de novo* sequencing.

**Conclusions:** This study provides a scalable framework for mutation profiling and discovery proteomics analysis that enables sensitive detection of both known and emerging viral variants in complex clinical samples.

## Background

Saliva is an attractive specimen for infectious disease diagnostics because it enables non-invasive sampling and captures diverse biological signals from the oral microenvironment. As a biologically informative biofluid, saliva provides a distinct window into human health. It contains host-derived molecules, resident microbiota, dietary components, and environmental inputs that together reflect host physiology, microbial ecology, and pathogen burden. Salivary metaproteomics has demonstrated that saliva is an ecosystem-level matrix containing both host-and microbiome-derived molecular signatures, making it a useful readout of oral physiological and ecological states [1–5]. Viral infection perturbs this ecosystem by inducing host-response signatures and reshaping oral microbial communities, thereby positioning saliva as a clinically relevant matrix for both pathogen detection and microenvironmental characterization [6–10]. However, the dynamic nature of saliva makes clinical diagnostics inherently challenging, as the large number of community members and the incompletely defined community composition that varies within individuals.

Current saliva viral diagnostics rely mainly on immunoassays and nucleic acid amplification tests. Although these approaches are clinically powerful, they depend on predefined antigenic or genomic targets and therefore capture only a limited dimension of infectious biology. Their performance can also be undermined by rapid viral evolution, as sequence divergence may impair antigen recognition, bias viral load estimation, or cause false-negative molecular results when diagnostic target regions no longer match circulating strains [11–14]. Although untargeted whole-genome sequencing approaches such as metagenomic next-generation sequencing (mNGS) can support viral detection, they can be affected by amplification bias, and often require extensive bioinformatic analysis for genome assembly and interpretation [15, 16]. Additionally, detectable levels of RNA are known to persist after SARS-CoV-2 infection has subsided [17].

Mass spectrometry (MS)–based proteomics provides a complementary framework for viral detection by directly detecting the presence of viral proteins. Both targeted and untargeted MS-based proteomics approaches to virus detection have been developed. During the SARS-CoV-2 pandemic, there was renewed interest in viral detection using proteomics that focused mainly on the targeted detection of SARS-CoV-2 to improve the limits of detection (LODs) and reproducibility through the measurement of predefined proteotypic targets [18–24]. In contrast, untargeted metaproteomics workflows make it possible to detect emerging and novel viral peptides from both sequenced and unsequenced variants, and to detect multiple viruses simultaneously. Untargeted approaches can also support full metaproteomic analyses to capture the proteomic state of both the host and the microbiome. Direct detection of SARS-CoV-2 proteins together with host proteome alterations in saliva further illustrates the potential of MS-based proteomics to unify viral detection with host-response profiling in a single assay [7]. However, the main challenge with untargeted viral detection, and metaproteomics in general, is that increasing database size decreases sensitivity of database correlation searches. To combat the loss of sensitivity and increase in search time that occurs with large database searches, a number of ingenious strategies have been developed to reduce the database size while limiting the exclusion of proteins that may be present in the sample and important to the analysis. vPro-MS generated a reduced *in silico* peptide library spanning 331 human-pathogenic viruses in UniProtKB, comprising about 122,000 virus-specific peptides for broad untargeted viral surveillance [25]. The Armengaud group introduced a cascade proteotyping strategy that begins with the NCBInr database and progressively refines taxonomic specificity from broader assignments to highly specific strain-level. It achieved accurate identification of 23 viral species across 53 public datasets and demonstrated clinical applicability by detecting vaccinia virus in saliva [26].

A further refinement of viral detection in saliva is the detection of viral variants, which requires an untargeted approach. Variant detection poses a particular challenge as many respiratory viruses mutate very rapidly [27, 28], and an efficient method to incorporate these mutated sequences has not been established. Single-stranded RNA viruses, such as influenza and SARS-CoV-2, have higher mutation rates than double stranded RNA and DNA viruses. Herpes simplex virus (HSV) is an example of a DNA virus with a much lower mutation rate. Genomic surveillance studies have shown that recent sequence data are critical for tracking viral evolution: SARS-CoV-2 outbreak.info reports used time-resolved lineage and mutation data to monitor rapid variant shifts [29], and influenza A/H3N2 forecasting improved when the prediction window was shortened from 12 to 6 months [30]. In our previous study, attomole-level detection of variant-specific SARS-CoV-2 peptides showed that MS can sensitively resolve protein-level sequence variation as a complement to genome-based surveillance [31]. Rajoria *et al.* demonstrated that a restricted in-house SARS-CoV-2 mutant peptide database can support LC-MS/MS detection of variant-associated peptides, but the approach remained reliant on similarity matching with a threshold to the UniProt reference sequence to predefined variants [32]. Again, the key unresolved challenges are how to balance analytical sensitivity with database size and how to efficiently update the database with new viral variants. Increasing database comprehensiveness improves theoretical coverage of circulating and emerging variants, yet it also enlarges the search space, which reduces the sensitivity of the database search [33], decreasing overall peptide identification. This trade-off is particularly acute in saliva, where viral proteins are often present at low abundance in a complex background dominated by the host and non-viral members of the microbiome.

In this study, we develop a strategy for saliva-based viral proteomics that addresses this trade-off by combining a tailored viral mutation database with a single amino acid substitution (SAAS) mutation discovery approach. Protein sequences of new viral variants are downloaded from the GISAID website [34]. To prevent the database from becoming too large, decreasing sensitivity and excessively slowing the search, variant sequences are culled using prevalence- and time-based filters to retain representative circulating variants. To enable the discovery of new viral mutations, Global PTM Discovery (G-PTM-D) in MetaMorpheus [35] is used to identify spectra that have a high probability of arising from a SAAS. Discovered sequences inferred from these spectra are then incorporated into the database to enable ongoing mutation monitoring. The key advances of this work are: prevalence-based filtering combined with time filtering to reduce loss of sensitivity, real-world sequence-based validation of database representativeness and timeliness, evaluation of relative performance based on quantitative measurements, integration with G-PTM-D for SAAS discovery, and extension beyond SARS-CoV-2 to multiple viruses in saliva. This strategy enables sensitive detection of both known and emerging viral variants while maintaining database tractability in complex saliva proteomes, while also complementing viral sequencing through direct protein-level characterization of sequence variation, including emerging mutations not represented in predefined reference databases.

## Results

### Overview of the mutation profiling and discovery proteomics strategy for virus detection in saliva

Extensive and rapidly accumulating viral mutations can substantially expand the sequence search space. Meanwhile, the low relative abundance of viral proteins in an intrinsically complex saliva background presents significant analytical challenges for sensitive proteomic studies. Together, these two factors create a fundamental trade-off as the sensitivity of protein identification by database correlation decreases with increasing database size [36, 37]. To overcome this challenge and evaluate the feasibility of MS-based proteomics for viral detection in saliva in a clinical diagnostic context, we developed a strategy that integrates a tailored viral mutation database with a mutation discovery approach, enabling detection of emerging viral variants while preserving high database-search sensitivity. Databases for the high-priority viruses, including SARS-CoV-2 and influenza (H1N1, H3N2, and influenza B), are publicly available from large sequencing efforts such as GISAID. For SARS-CoV-2, H1N1, H3N2, and influenza B, both genomic and proteomic databases are available for variants of concern and variants of interest. However, downloading all of the variant sequences was over 100 GB as of March 18^th^, 2026, creating the need for more efficient database searching approaches. Thus, we have developed a strategy that greatly shrinks database size while retaining sequences of higher clinical relevance. At the same time, our strategy preserves the discovery capability of detecting single amino acid mutations, enabling the detection of emerging and previously unannotated viral mutations beyond predefined reference sequences.

Not all sequence variation from the large sequencing efforts is equally informative for clinical diagnostics, and much of it introduces unnecessary redundancy into the search database. In particular, outdated sequences are unlikely to reflect currently circulating variants, whereas low-prevalence mutations contribute limited diagnostic value. Accordingly, we generated a viral mutation peptide database using temporal and prevalence filters to optimize the balance between database coverage and search sensitivity (Fig. 1 Module 1). The 3-month time window was designed to prioritize recently circulating variants and capture mutations most likely to be detected in contemporary samples. It reduced database complexity by 99.8% for SARS-CoV-2, retaining 33,184 of 17,462,874 candidate sequences, and by 89.3% for influenza (H1N1, H3N2, and influenza B), retaining 55,489 of 519,773 candidate sequenced viruses. All reported mutated peptides were then aggregated at the lineage level, and a frequency threshold was applied to limit the candidate search space while preserving prevalent and representative viral variation. Each peptide was annotated with its protein position, viral lineage, clade, and collection date for downstream analysis.

**Figure 1:**
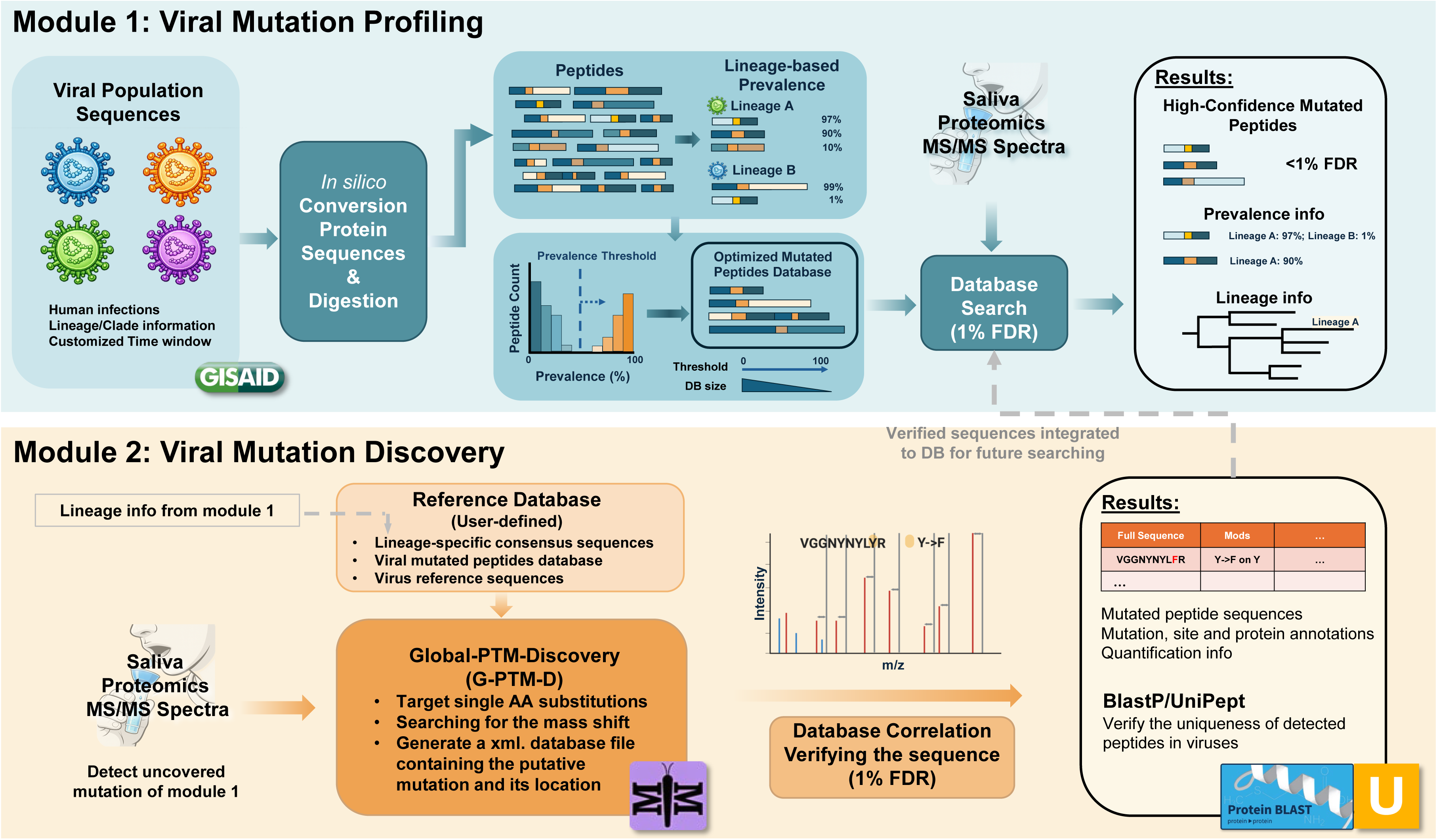
Schematic overview of mutation profiling and discovery proteomics strategy. To detect multiple viruses, known viral variants, and new variants, both modules are used in parallel. To reduce the loss of sensitivity that results from adding all new viral sequences from genomic viral surveillance programs, such as GISAID, to the database, an *in silico* method for retaining the most relevant sequences by using both prevalence and temporal filters is presented (Module 1). The prevalence or occupancy of any mutation can vary from <1% to 100% within a viral variant. A tailored database is created from the reference and variant sequences from the *in silico* analyses, and database correlation search is performed on this tailored database. Peptide identifications are filtered at 1% FDR to obtain high-confidence mutated peptides and lineage-associated results. In Module 2, a type of quasi-open search that detects mass shifts corresponding to single amino acid mutations is performed in MetaMorpheus using the Global PTM Discovery function. The user-defined reference database for mutation discovery is the same database that is created in Module 1. Verified peptide identifications from mutation discovery can be added to the database for future detection.

To enable the discovery of emerging viral mutations, we leveraged the G-PTM-D feature within MetaMorpheus (Fig. 1 Module 2). This approach interrogates spectra for mass shifts consistent with SAASs by performing a quasi-open search against a reference database, which can be defined either from the lineage information generated in the previous step or according to user-specified criteria. Mutation candidates discovered through this process are then systematically integrated into a mutation-inclusive database, facilitating efficient downstream database correlation analyses. A following peptide verification is conducted by database correlation with FDR control, and sequence similarity analysis by Unipept [38] or Blast-P [39] is required to confirm that the mutated peptide is unique to the putative viral lineage. This strategy improves detection sensitivity for novel viral peptides while mitigating the increased false discovery rate (FDR) that accompanies iterative or excessively broad search workflows. Moreover, because database expansion is guided by observed data rather than exhaustive enumeration, the strategy remains computationally tractable and requires only limited computational resources for the viral targets under investigation.

### Prevalence thresholding reduces the mutated peptide database size by excluding rare mutations

Previous studies have shown that viral evolution continually generates large numbers of low-frequency mutations, whereas natural selection drives only a limited subset of advantageous variants to high prevalence [40, 41]. Consistent with this principle, our analysis of SARS-CoV-2 mutations from GISAID revealed a strongly bimodal prevalence distribution, with most mutations occurring either at very high prevalence (>90%) or very low prevalence (<1%) (Additional file 1: Fig. S1). Importantly, this feature was also observed at the peptide level. The majority of mutated peptides (43,632) fell into the >90% prevalence group for SARS-CoV-2 followed by 36,062 mutated peptides observed <5% prevalence (Fig. 2A). Influenza viruses: H1N1, H3N2, and influenza B also showed bimodal distributions, but with the opposite trend: most mutated peptides has less than 5% prevalence (Fig. 2B-D), consistent with the rapid evolution of influenza viruses. This bimodal property provides a practical basis for reducing database size while preserving representative coverage of circulating viral variants. We therefore generated peptide databases using lineage-level mutation prevalence thresholds of 0%, 5%, 10%, 20%, 30%, 40%, 50%, 60%, 70%, 80%, and 90% for SARS-CoV-2 and combined influenza (H1N1, H3N2, and influenza B) viruses, respectively, and evaluated their impact on search sensitivity.

**Figure 2:**
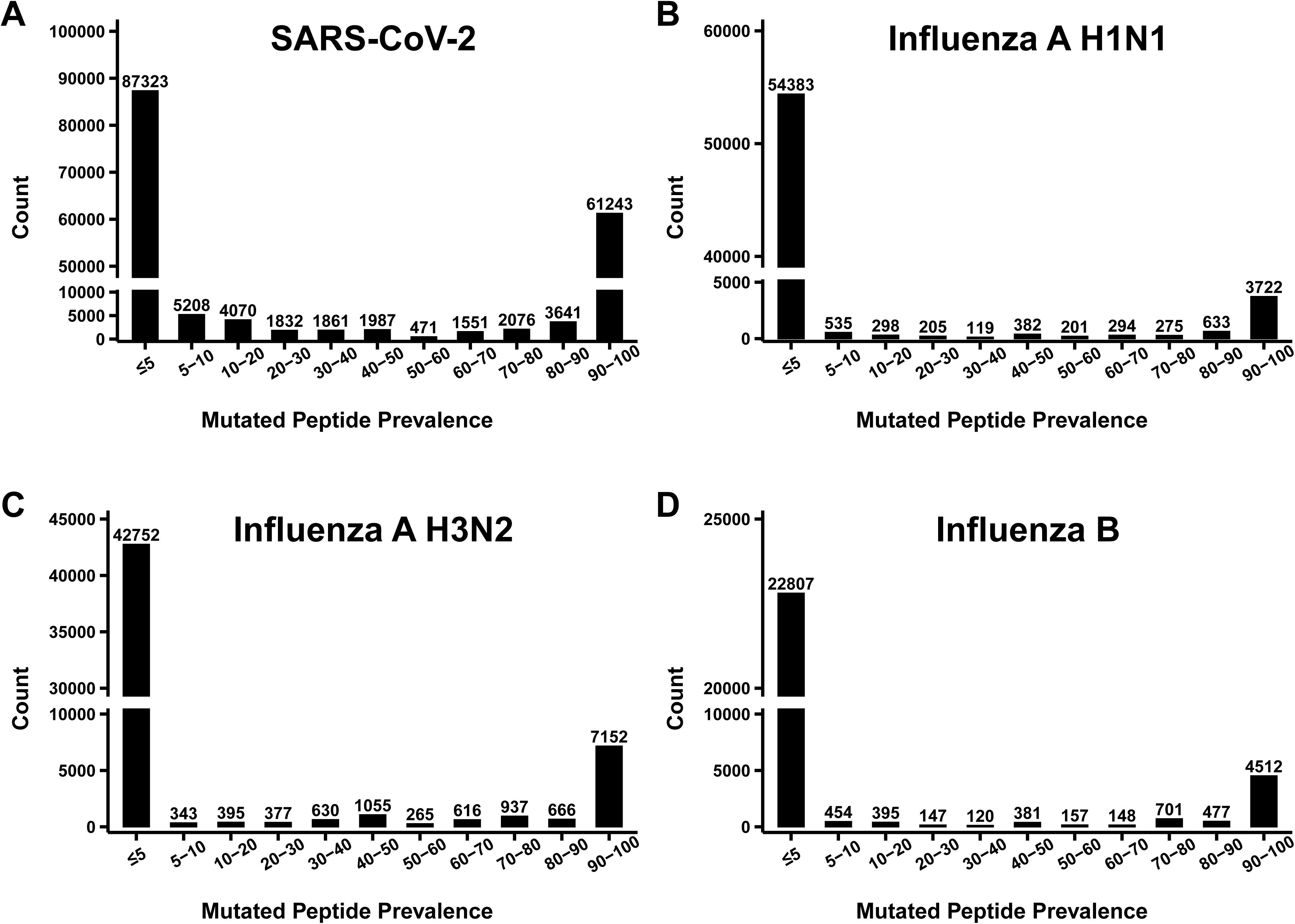
Bimodal prevalence distributions of mutated peptides across respiratory viruses. The prevalence thresholds are used to remove the large number of sequences with mutations that are rarely observed within a viral lineage. A, Prevalence distribution of SARS-CoV-2 mutated peptides from viral population sequences collected between June 1 and August 31, 2025, showing a bimodal pattern with the largest number of peptides in the >90% prevalence group (43,632 peptides), relatively few peptides between 10% and 90% (17,489 peptides), followed by the <5% prevalence group (36,062 peptides). B–D, Prevalence distributions of mutated peptides from influenza A/H1N1, influenza A/H3N2, and influenza B, based on viral population sequences collected between November 1, 2024, and February 28, 2025, also showed bimodal patterns, with the majority of mutated peptides occurring in the low-prevalence range (<5%).

Thus, we further applied prevalence thresholds and evaluated reductions in mutated peptide database size under low- and high-prevalence filtering criteria. For SARS-CoV-2, the database decreased from 39,644 peptides at the 0% threshold to 8,859 peptides at the 5% threshold, corresponding to a 77.7% reduction, and to 7,090 peptides at the 10% threshold, corresponding to an 82.1% reduction (Fig. 3A). Increasing the threshold from 10% to 90% further reduced the database to 2,875 peptides, equivalent to an additional 10.6% reduction relative to the original 0% database. For influenza, the database decreased from 107,159 peptides at the 0% threshold to 6,696 peptides at the 5% threshold, corresponding to a 93.8% reduction, and to 5,831 peptides at the 10% threshold, corresponding to a 94.6% reduction (Fig. 3B). Increasing the threshold from 10% to 90% further reduced the database to 3,970 peptides, equivalent to an additional 1.7% reduction relative to the original 0% database. Because a larger database increases the multiple-hypothesis burden and intensifies penalties in target–decoy competition (TDC) [42], these reductions are expected to improve sensitivity at a fixed FDR by lowering the score threshold required for acceptance. In addition, the smaller databases can be searched much more rapidly and decrease the need for sophisticated computer hardware.

**Figure 3:**
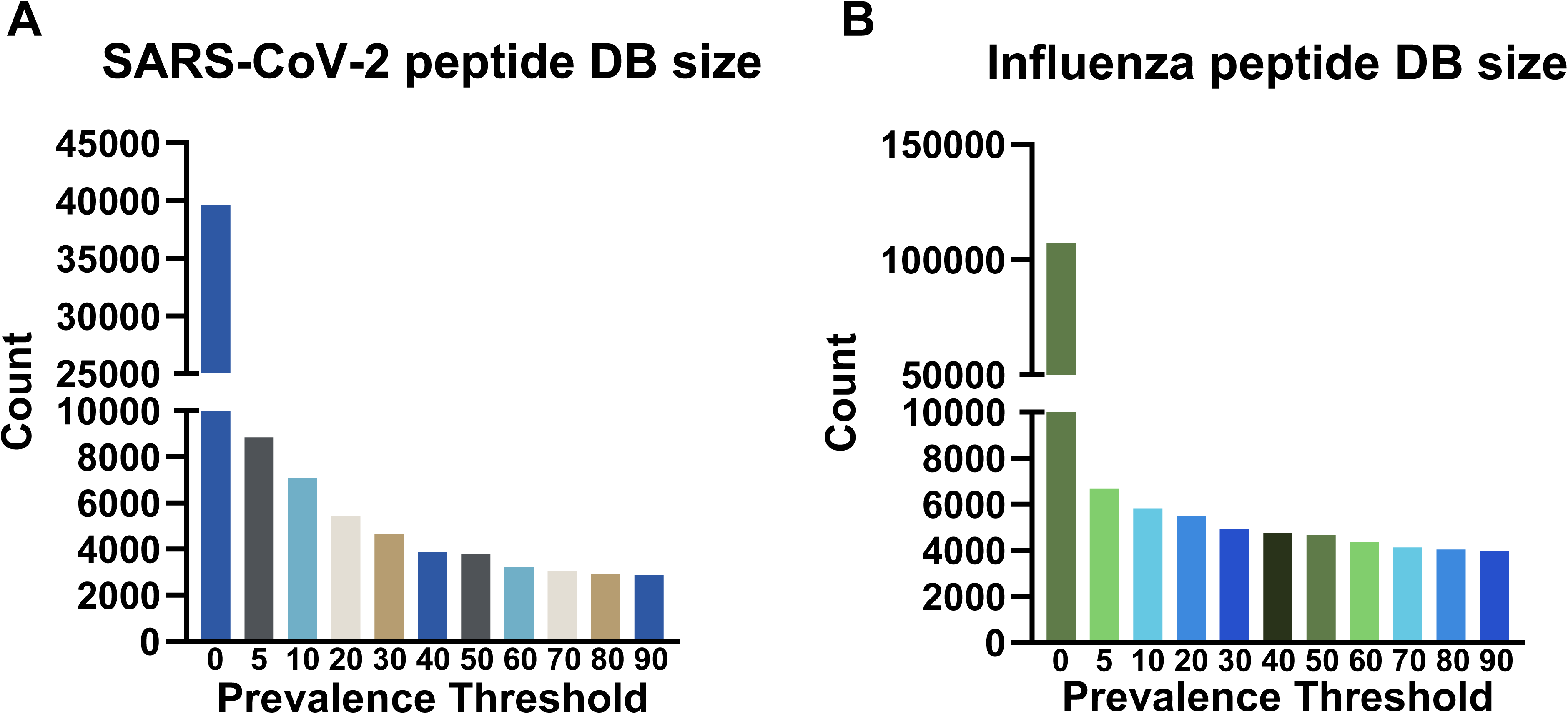
Prevalence based thresholding substantially reduces mutated peptide database size. A, SARS-CoV-2 mutated peptide database size across prevalence thresholds. Database size decreased by 77.7% at the 5% threshold and by 82.1% at the 10% threshold, with a further reduction of 10.6% at the 90% threshold relative to the original database. B, Influenza mutated peptide database size across prevalence thresholds. Database size decreased by 93.8% at the 5% threshold and by 94.6% at the 10% threshold, with a further reduction of 1.7% at the 90% threshold relative to the original database.

### Tailored mutation peptide database reduces sensitivity loss in database search

To quantify the trade-off between database size and search sensitivity, we analyzed cultured SARS-CoV-2 and H1N1 samples against mutation peptide databases generated using prevalence thresholds to limit the increase in database size. Both viruses were analyzed in their original culture matrices, where viral proteins constituted only a minor fraction of the total sample protein content. The culled mutated sequences were combined with the corresponding viral reference sequences and contaminants including proteins from host cells and culture background. Compared with the control search containing only viral reference sequences and contaminants, lower prevalence thresholds led to progressive reductions in peptide-spectrum match (PSM) identifications as the mutation database increased in size. The greatest loss was observed when no prevalence threshold was applied, resulting in an average loss of 306 PSMs for SARS-CoV-2 (Fig. 4A) and 299 PSMs for influenza H1N1 (Fig. 4B) relative to the control searches, corresponding to 4.1% and 11.0% of control PSMs, respectively. Applying a 5% prevalence threshold effectively controlled sensitivity loss, keeping it below 1% in both cases. A smaller but measurable decline was also observed at the 90% threshold, indicating that even inclusion of only high-prevalence mutation peptides imposed a search penalty. Although the 5% threshold performed markedly better than the unfiltered database, PSM loss remained approximately 3-fold higher than that observed at the 90%, 80%, and 70% thresholds for SARS-CoV-2, and 1.7-fold higher for influenza.

**Figure 4:**
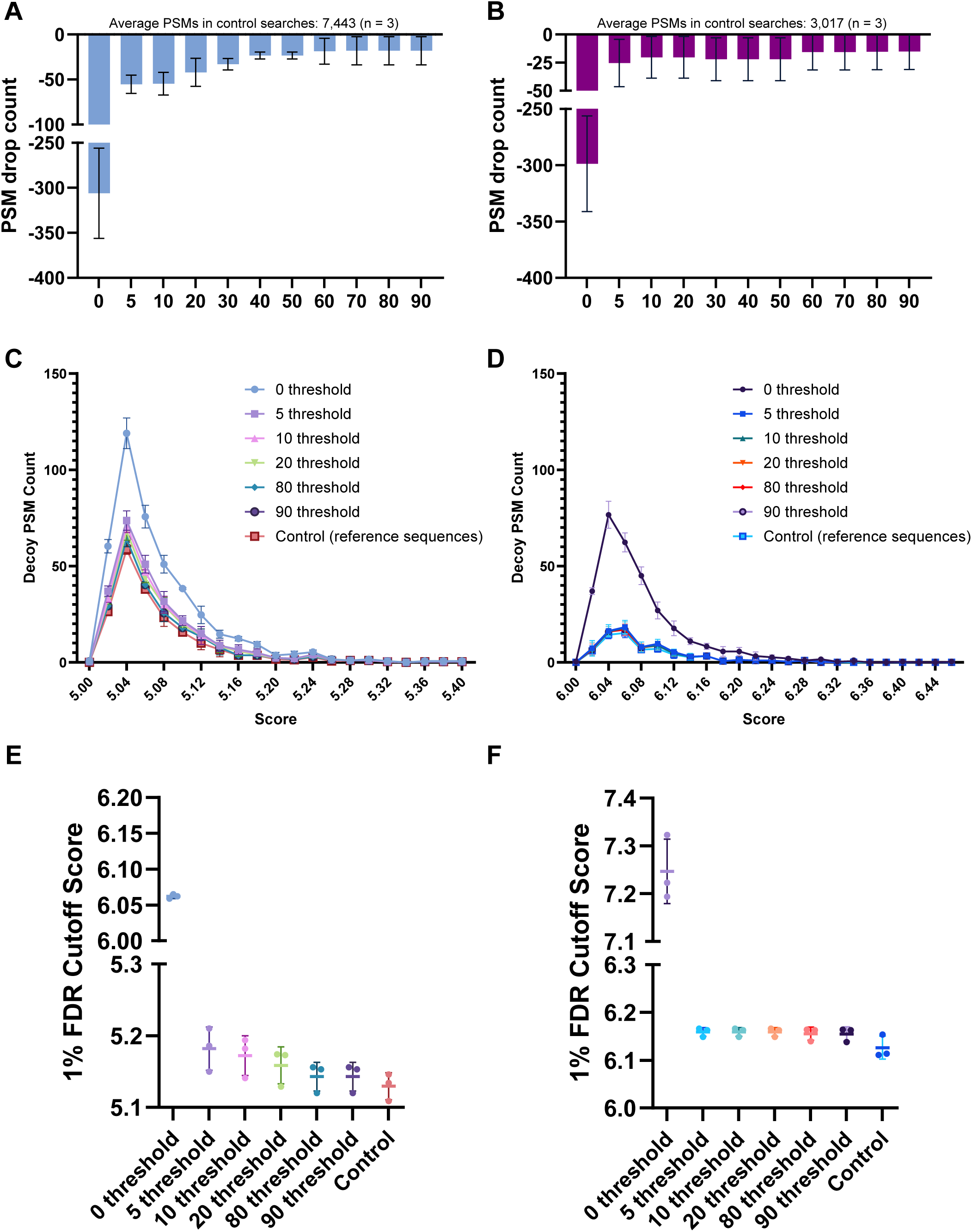
Prevalence threshold greatly reduces search sensitivity loss caused by database size inflation. A, B, Increasing the prevalence threshold increases the number of peptide spectrum matches (PSMs) from cultured SARS-CoV-2 (A) and influenza A H1N1 (B) samples compared to the viral reference sequences and contaminants. C, D, The number of PSMs matching decoy sequences increases with database size in SARS-CoV-2 (C) and influenza A H1N1 (D) searches, and the reduction in decoy matches with increasing prevalence threshold is shown. E, F, The PSM score cutoffs required to maintain 1% FDR control for SARS-CoV-2 (E) and influenza A/H1N1 (F) decrease with higher applied thresholds, showing the sensitivity enhancement with higher prevalence thresholds and smaller databases.

To further demonstrate how database expansion decreases search sensitivity, we interrogated peptide-spectrum match (PSM)-level behavior. Expansion of the target database proportionally increased the size of the decoy database, thereby elevating the probability of random matches during TDC. Accordingly, searches against the unfiltered mutated peptide databases produced substantially more decoy PSMs for both SARS-CoV-2 and influenza A H1N1 (Fig. 4C, D). Together, these findings indicate that prevalence thresholding should be optimized according to the viral lineage and the analytical objective, with the associated sensitivity trade-off consistently observed across search engines, including FragPipe [43] (Additional file 1: Fig. S2), and in virus-spiked saliva samples (Additional file 1: Fig. S3).

### Tailored database maintains the breadth of search for SARS-CoV-2 and influenza viral mutations

Applying lineage-level mutation prevalence thresholds improves database-search sensitivity, while retaining the most clinically relevant sequences. Prevalence filtering enriches for mutations above a defined within-lineage frequency; however, abrupt shifts in mutation prevalence (e.g., rapid expansion of a newly emerging variant) and various stochastic mutations can violate this assumption and reduce coverage. Consequently, the high mutation rates of SARS-CoV-2 and influenza necessitate explicit evaluation of whether a threshold database remains sufficiently comprehensive for upcoming viral population detection.

To determine the coverage and test the timeliness of our strategy, we used sequences from the subsequent month in GISAID for both SARS-CoV-2 and influenza (H1N1, H3N2, and influenza B) and quantified the fraction of mutated peptides in the new-month population covered by databases generated at different thresholds. For all threshold settings, overall coverage remained high and exceeded 94% across conditions. The 0% prevalence-threshold databases captured 99.59% of the SARS-CoV-2 viral population (Fig. 5A) and 99.68% of the influenza viral population (Fig. 5B), demonstrating the feasibility of retaining near-complete coverage of forthcoming mutational diversity. A low 5% threshold databases maintained 98.2% for SARS-CoV-2 and 98.5% for influenza, while significantly decreasing the database size. For SARS-CoV-2, raising the prevalence threshold from 20% to 80% caused only a modest reduction in coverage (0.8%). In contrast, the same threshold change resulted in a 2.4% loss of coverage for influenza, indicating that threshold selection is also virus dependent. These results indicate that moderate prevalence filtering with an appropriate time window can yield a compact database that remains highly representative of near-future circulating virus diversity.

**Figure 5:**
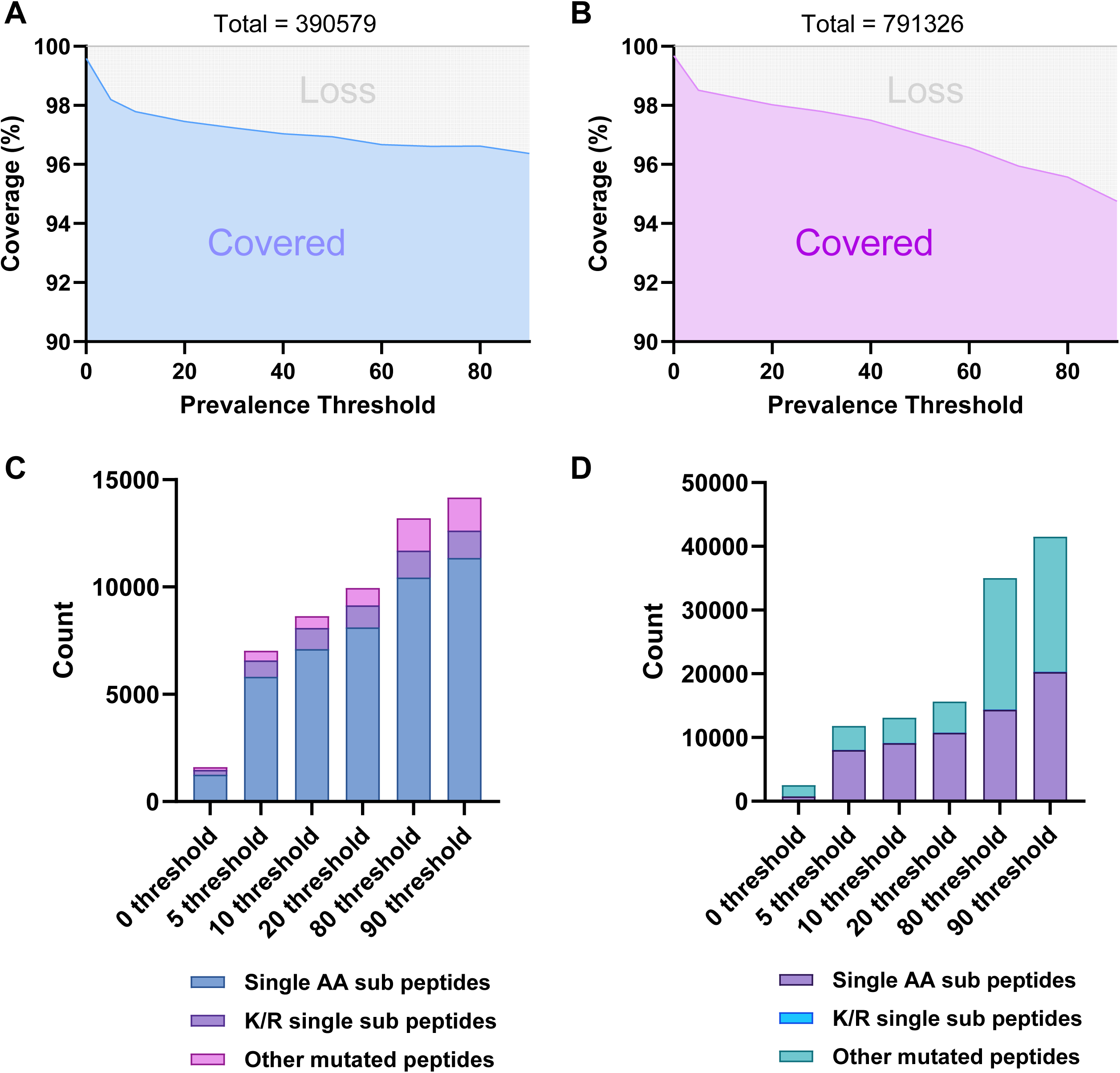
Prevalence-filtered mutation peptide databases retain broad coverage of near-future viral diversity. A, B, Fraction of mutated peptides in subsequent-month GISAID viral populations covered by databases generated at different prevalence thresholds for SARS-CoV-2 (A) and influenza (B; H1N1, H3N2, and influenza B). Coverage remained high across all threshold settings and exceeded 94% in all conditions. The uncovered peptides have the lowest prevalence or site occupancy within the reported sequences for the viral variant, and presumably have reduced clinical relevance. C, D, Composition of peptides not covered by prevalence-filtered databases at different thresholds for SARS-CoV-2 (C) and influenza (D). Most uncovered peptides were single amino acid substitution peptides. K/R substitutions are shown separately because they disrupt tryptic cleavage and require separate handling.

To further demonstrate the effectiveness of this strategy in preserving informative viral mutations, we examined rapidly evolving antigenic proteins, including the SARS-CoV-2 spike protein and influenza hemagglutinin (HA) and neuraminidase (NA). For SARS-CoV-2 spike protein mutations, all thresholds maintained near-complete coverage with the lowest coverage of 97.9% at the 90% prevalence threshold (Table 1). In contrast, influenza exhibited a stronger trade-off at higher thresholds: coverage was 98.0% at a 20% prevalence threshold for HA (Table 2) and 94.5% for NA (Table 3), whereas coverage dropped to 89.1% for HA and 89.8% for NA at an 80% prevalence threshold. These results indicate that the optimal prevalence threshold should be selected according to viral evolutionary dynamics. Thus, influenza database construction requires an intermediate prevalence threshold (<20%) to preserve high representativeness, whereas a more stringent threshold works better for SARS-CoV-2.

**Table 1.** Prevalence-threshold effects on SARS-CoV-2 spike mutation coverage. Coverage of mutated peptides from the SARS-CoV-2 spike protein by databases constructed using different prevalence thresholds. Near-complete coverage was retained across all thresholds, with the lowest value of 97.9% observed at the 90% threshold.

| Coverage of SARS-CoV-2 Sep 2025 Spike protein |  |  |  |  |  |  |
| --- | --- | --- | --- | --- | --- | --- |
|  | 0 threshold | 5 threshold | 10 threshold | 20 threshold | 80 threshold | 90 threshold |
| Exact Matched Peptides | 171260 (99.9%) | 170412 (99.4%) | 170037 (99.2%) | 169711 (99.0%) | 168410 (98.2%) | 167951 (97.9%) |
| Single AA sub peptides | 177 (0.0%) | 804 (0.5%) | 993 (0.6%) | 1138 (0.7%) | 1818 (1.1%) | 2233 (1.3%) |
| K/R single sub peptides | 20 (0.0%) | 94 (0.1%) | 240 (0.1%) | 247 (0.1%) | 265 (0.2%) | 277 (0.2%) |
| Other mutated peptides | 27 (0.0%) | 174 (0.1%) | 214 (0.1%) | 388 (0.2%) | 991 (0.6%) | 1023 (0.6%) |

**Table 2.** Prevalence-threshold effects on influenza HA mutation coverage. Coverage of mutated peptides from influenza hemagglutinin (HA) by databases constructed using different prevalence thresholds. Intermediate filtering retained high coverage (98.0% at 20%), whereas stringent filtering reduced coverage to 89.1% at 80%.

| Coverage of Influenza March 2025 HA |  |  |  |  |  |  |
| --- | --- | --- | --- | --- | --- | --- |
|  | 0 threshold | 5 threshold | 10 threshold | 20 threshold | 80 threshold | 90 threshold |
| Exact Matched Peptides | 87602 (99.7%) | 86609 (98.6%) | 86130 (98.0%) | 86127 (98.0%) | 78320 (89.1%) | 78320 (89.1%) |
| Single AA sub peptides | 82 (0.1%) | 855 (1.0%) | 1163 (1.3%) | 1166 (1.3%) | 1070 (1.2%) | 1070 (1.2%) |
| K/R single sub peptides | 2 (0.0%) | 4 (0.0%) | 4 (0.0%) | 4 (0.0%) | 2 (0.0%) | 2 (0.0%) |
| Other mutated peptides | 186 (0.2%) | 404 (0.5%) | 575 (0.7%) | 575 (0.7%) | 8480 (9.7%) | 8480 (9.7%) |

**Table 3.** Prevalence-threshold effects on influenza NA mutation coverage. Coverage of mutated peptides from influenza neuraminidase (NA) by databases constructed using different prevalence thresholds. Coverage was maintained at 94.5% at the 20% threshold but decreased to 89.8% at the 80% threshold.

| Coverage of Influenza March 2025 NA |  |  |  |  |  |  |
| --- | --- | --- | --- | --- | --- | --- |
|  | 0 threshold | 5 threshold | 10 threshold | 20 threshold | 80 threshold | 90 threshold |
| Exact Matched Peptides | 63816 (99.6%) | 61797 (96.5%) | 61746 (96.4%) | 60544 (94.5%) | 57520 (89.8%) | 57520 (89.8%) |
| Single AA sub peptides | 126 (0.2%) | 1368 (2.1%) | 1411 (2.2%) | 1775 (2.8%) | 3002 (4.7%) | 3002 (4.7%) |
| K/R single sub peptides | 0 (0.0%) | 0 (0.0%) | 0 (0.0%) | 0 (0.0%) | 0 (0.0%) | 0 (0.0%) |
| Other mutated peptides | 116 (0.2%) | 893 (1.4%) | 901 (1.4%) | 1739 (2.7%) | 3536 (5.5%) | 3536 (5.5%) |

Notably, among peptides not covered by the intermediate prevalence (<20%) threshold databases, most differed by only a single amino acid substitution from a database peptide (81.5% for SARS-CoV-2 (Fig. 5C); and 68.4% for influenza (Fig. 5D) at 20% prevalence threshold). This indicates that SAAS are the dominant class of residual uncovered variation after prevalence filtering and can be recovered by G-PTM-D through detection of the corresponding substituted amino acid residue mass shifts.

### Determining the LODs of viral peptides and lysate in saliva for SARS-CoV-2, influenza A, H1N1, and HSV-1

To mimic realistic infection scenarios, we further performed a metaproteomics analysis on detecting multiple viruses within saliva samples to determine the overall sensitivity of our method. A 4 μL aliquot of SARS-CoV-2 viral preparation containing viral material, host-cell components, and culture-medium background was spiked into 80 μL of saliva and designated as the 1× input. The viral preparation contributed 3.56 μg total protein. Additional virus spike-in dilutions of 0.5×, 0.1×, 0.05×, and 0.01× into 80 μL saliva were analyzed to determine the LODs. For database searching, a 10% prevalence threshold was applied during database optimization for SARS-CoV-2, and influenza (H1N1, H3N2, and influenza B), and the resulting viral peptide databases were combined for simultaneous detection of multiple viruses in each sample. An in-house salivary protein database was included as the target database for saliva metaproteomics analysis. Using this strategy, we identified and quantified 30 SARS-CoV-2 tryptic peptides together with 1,631 proteins derived from human, bacterial, and other background sources in the 1× diluted sample. Nucleocapsid phosphoprotein was the dominant viral protein, represented by 19 peptides, and 2 nucleocapsid peptides were consistently detected and quantified at the lowest detected concentration in the 0.05× diluted sample (Fig. 6A), highlighting their potential as biomarkers for viral detection.

**Figure 6:**
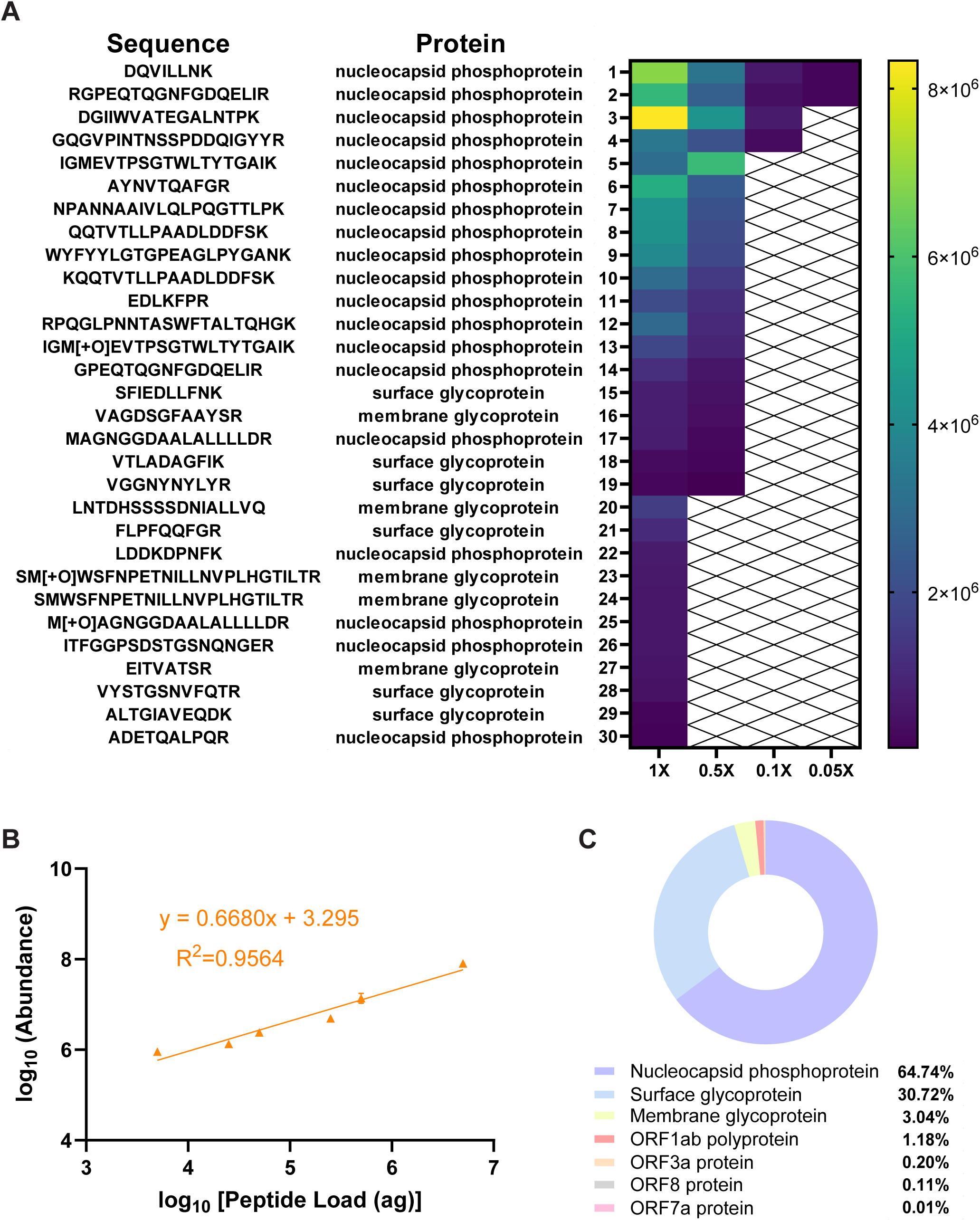
LOD determination for SARS-CoV-2 in saliva using 10% prevalence threshold. A, Heat map of SARS-CoV-2 peptides detected across a saliva spike-in dilution series (1×, 0.5×, 0.1×, and 0.05×). A total of 30 tryptic peptides were identified, including 19 from nucleocapsid phosphoprotein, with two nucleocapsid peptides detected at the lowest concentration (0.05×). B, Log-log calibration curve for the reference peptide VGGNYNYLYR in saliva matrix (R² = 0.9564). C, Mass fractions of SARS-CoV-2 proteins determined by the total protein approach, with nucleocapsid phosphoprotein and spike glycoprotein comprising 73.65% and 22.26% of total protein mass, respectively. The estimated limit of detection was 9.4 ng viral lysate per mL saliva.

For absolute viral peptide quantification, the reference peptide VGGNYNYLYR was spiked into a saliva matrix as a dilution series for LC-MS/MS analysis. A calibration curve was generated by a log–log linear regression model to minimize the bias from exponential dilution. The result demonstrated a strong quantitative correlation (R² = 0.9564) (Fig. 6B), enabling precise conversion from MS peptide intensity to actual protein amount. Next, viral protein composition profiling was conducted by analyzing viral lysate only. The total protein approach (TPA)[44] further revealed nucleocapsid phosphoprotein as the predominant component (73.65% of total protein mass), followed by spike glycoprotein (22.26%) (Fig. 6C). We then used this quantitative measure with the diluted virus-spiked saliva dilution series to convert from peptide-level quantification into viral mass equivalents. From this analysis, we determined that our proteomics strategy can reliably detect as little as 9.4 ng of viral lysate per mL of saliva. Based on the average estimate of 48 spike trimers per SARS-CoV-2 virion by Klein et al. [45], the measured LOD corresponds to approximately 3.4 × 10^4^ virions per mL of saliva. This LOD is similar to results obtained for other viral proteins, in which orbitrap of similar MS systems were used [31].

Similar analyses were carried out for influenza H1N1 and HSV-1. HSV-1 was included as a common and highly transmissible DNA virus with a more diverse proteome (with about 80 proteins) that does not rapidly mutate. The resulting LODs were 58.4 ng/mL saliva for H1N1 (Fig. 7) and 1.29 μg/mL saliva for HSV-1, respectively (Additional file 1: Fig. S4). Applying the corresponding conversion to influenza H1N1 (375 HA trimers per virion) [46] and HSV-1 (300 glycoprotein D per virion) [47, 48] yielded LODs of 1.27 × 10^5^ virions/mL saliva and 1.82 × 10^5^ virions/mL saliva, respectively. These results confirmed that our mutation profiling proteomics strategy is broadly applicable for the detection and quantification of various viral pathogens in saliva. Its high sensitivity, combined with precise annotation and robustness against complex biological backgrounds, underscores its potential for clinical diagnostic applications and large-scale surveillance of viral infections.

**Figure 7:**
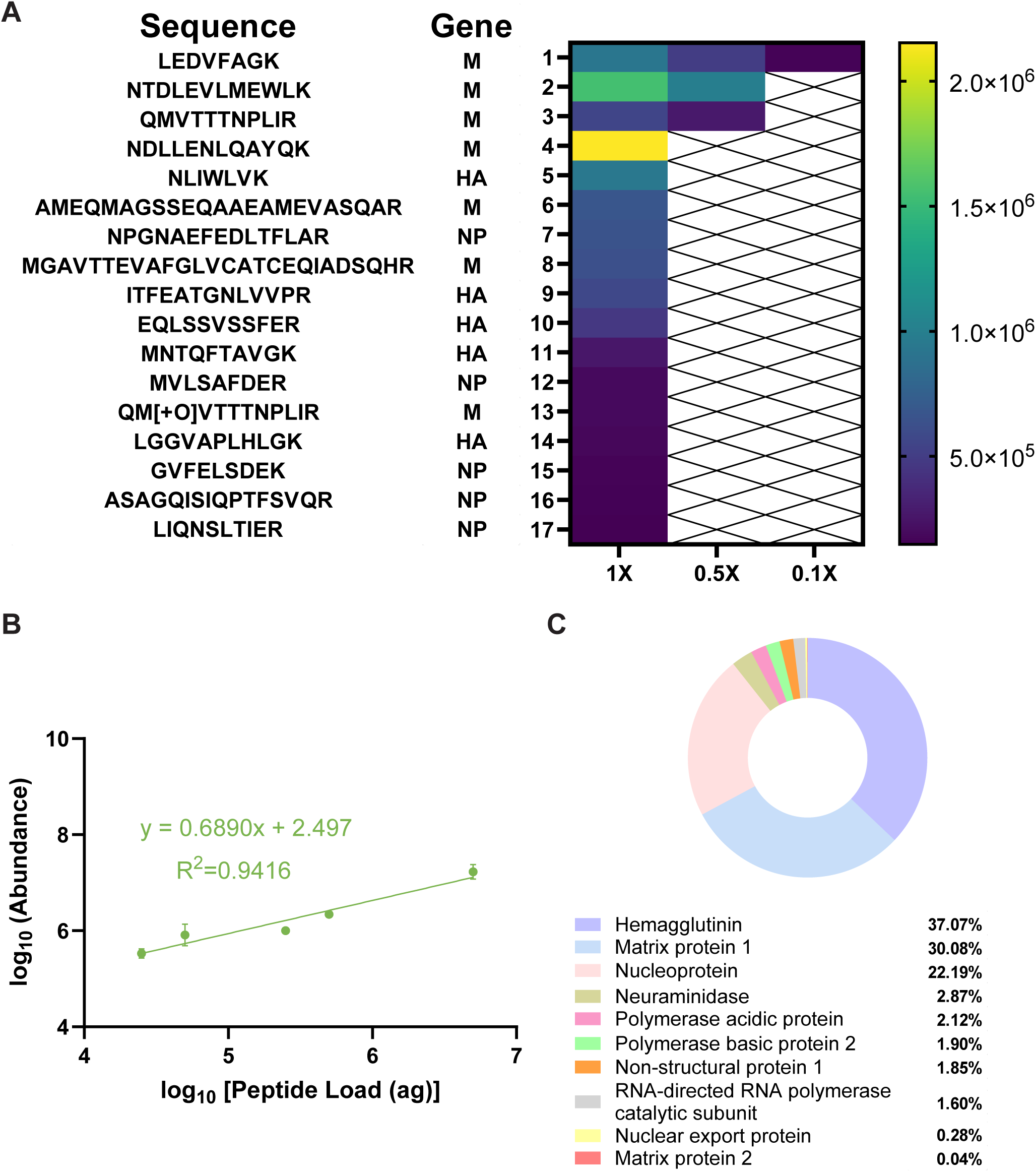
LOD determination for Influenza A H1N1 in saliva using 10% prevalence threshold. A, Heat map of influenza A H1N1 peptides detected across a saliva spike-in dilution series (1×, 0.5×, and 0.1×). A total of 17 tryptic peptides were identified, including 7 from matrix protein 1, with 1 nucleocapsid peptides detected at the lowest concentration (0.1×). B, Log-log calibration curve for the reference peptide LGGVAPLHLGK in saliva matrix (R² = 0.9416). C, Mass fractions of influenza A H1N1 proteins determined by the total protein approach. The estimated limit of detection was 58.4 ng viral lysate per mL saliva.

### G-PTM-D detects unknown single AA variants with better sensitivity than conventional PTM search and *de novo* sequencing

To detect new sequences not found in databases, we utilized the G-PTM-D feature in MetaMorpheus. A quasi-open search is performed that searches for the mass shift of single AA mutations over the peptide of interest [35, 49]. All possible SAASs are activated. A *.xml file is generated containing the putative mutation and its location which can later serve as a mutation database of the regular database search task for final confirmation with FDR control. In principle, this strategy can recover the majority of peptides excluded by the prevalence-threshold filter, as described in the previous section.

To demonstrate the sensitivity of our strategy for detection of unknown viral mutations, we selected the spike protein from the SARS-CoV-2 virus as our target of interest. Next, we spiked two mutated peptides, VGGNYNYLFR (Y453F) and NTQQVFAQVK (E780Q), along with one unmutated reference peptide, VGGNYNYLYR, into saliva samples and made a dilution series. The Y453F mutation has previously been associated with antigenic drift and host adaptation [50, 51], and E780Q is in the S2 subunit which mediates viral cell membrane fusion [52]. VGGNYNYLYR and NTQEVFAQVK represent the original peptide sequences of the spike protein in the reference database for the G-PTM-D search. As a result, both reference and mutated peptides were detected at loaded peptide amounts as low as 50 fg, with PSM counts reported in Table 4. These LODs correspond to 41.6 amol for VGGNYNYLFR and 43.0 amol for NTQQVFAQVK. Both mutated peptides were identified and quantified with proper annotation of the original sequences, illustrating the ability to concurrently detect multiple sequence variants. Moreover, final database correlation after G-PTM-D showed the same PSM counts as direct searching with a customized mutation-inclusive database across all peptide loads, indicating that the modest, targeted database expansion introduced by G-PTM-D did not reduce search sensitivity (Additional file 2: Table S1).

**Table 4.** Detection of reference and mutated SARS-CoV-2 spike peptides by G-PTM-D in saliva background. PSM counts for the identified reference peptide VGGNYNYLYR and the mutated peptides VGGNYNYLF[Y453F]R and NTQQ[E780Q]VFAQVK in saliva peptide background across a dilution series. G-PTM-D in MetaMorpheus identified both reference and mutated peptides at concentrations as low as 50 fg, with similar detection performance for mutated and reference sequences.

| <u>Viral Mutation Discovery</u> | VGGNYNYLYR (ref.) |  | VGGNYNYLF[Y453F]R |  | NTQQ[E780Q]VFAQVK |  |
| --- | --- | --- | --- | --- | --- | --- |
| Peptide load (fg) | Rep. 1 | Rep. 2 | Rep. 1 | Rep. 2 | Rep. 1 | Rep. 2 |
| 2500 | <u>2 PSMs</u> | <u>2 PSMs</u> | <u>1 PSM</u> | <u>1 PSM</u> | <u>1 PSM</u> | <u>1 PSM</u> |
| 500 | <u>1 PSM</u> | <u>2 PSMs</u> | <u>1 PSM</u> | <u>1 PSM</u> | <u>1 PSM</u> | <u>1 PSM</u> |
| 250 | <u>2 PSMs</u> | <u>1 PSM</u> | <u>1 PSM</u> | <u>1 PSM</u> | <u>1 PSM</u> | <u>1 PSM</u> |
| 50 | <u>2 PSMs</u> | <u>2 PSMs</u> | <u>1 PSM</u> | <u>1 PSM</u> | <u>1 PSM</u> | <u>1 PSM</u> |
| 10 | 0 PSM | 0 PSM | 0 PSM | 0 PSM | 0 PSM | 0 PSM |

Comparing G-PTM-D with a conventional PTM search for SAAS revealed that G-PTM-D preserved search sensitivity, whereas the expanded search space of conventional PTM searching led to substantial identification loss. Conventional PTM searches substantially increase *in silico* database entropy, which reduces sensitivity and often misses low-abundance peptides. In our case, the conventional PTM approach failed to detect any spiked-in reference or SAAV peptides, while also causing an approximately 52% reduction in identified protein groups (Table 5). Conventional PTM searches not only increase computational time but also intensify FDR penalties, because the decoy search space expands simultaneously as multiple potential PTMs are considered.

**Table 5.** Detection of reference and mutated SARS-CoV-2 spike peptides by conventional PTM search in saliva background. Comparison of protein group identifications obtained using the optimized G-PTM-D-based strategy and a conventional PTM search. The conventional PTM search failed to detect the spiked reference or mutated peptides and reduced the number of identified protein groups by approximately 52%.

| <u>Conventional PTM search</u><br><u>with single AA substitutions</u> | VGGNYNYLYR (ref.) | VGGNYNYLF[Y453F]R | NTQQ[E780Q]VFAQVK | ProteinID Drop<br>% |
| --- | --- | --- | --- | --- |
| Peptide load (fg) | Rep. 1 &2 | Rep. 1 &2 | Rep. 1 &2 | Compared to No PTM search |
| 2500 | Fail | Fail | Fail | 48.3% |
| 500 | Fail | Fail | Fail | 57.1% |
| 250 | Fail | Fail | Fail | 42.9% |
| 50 | Fail | Fail | Fail | 59.6% |

Moreover, *de novo* sequencing provides a possible alternative for mutated peptide discovery because it does not require prior inclusion of the mutated sequence in a database, thereby reducing potential database-derived bias between reference and mutated peptides. Its performance, however, depends on MS/MS spectral quality and requires comprehensive fragmentation, which can be influenced by peptide sequence, abundance, and instrument settings. The decision-tree-based *de novo* sequencing tool Novor [53] performed well for the mutated peptide VGGNYNYLFR, correctly sequencing it across all tested conditions with PSM counts comparable to G-PTM-D (Fig. 8). For the other peptides, Novor showed less consistent performance, with only partial recovery of the correct sequence across peptide loads of mutated peptide NTQQVFAQVK. An abundance-dependent effect was also observed for the reference peptide VGGNYNYLYR, for which Novor correctly reported only a subset of PSMs at peptide loads ≥500 fg and did not recover the correct sequence below 500 fg. In contrast, the transformer-based *de novo* sequencing tool Casanovo [54] achieved better performance, correctly sequencing all reported PSMs for both mutated peptides similarly to the G-PTM-D approach. At the same time, Casanovo recovered the reference sequence across all peptide loads, although several predictions had low confidence scores, including 2 of 4 PSMs at 2500 fg, 2 of 3 PSMs at 500 fg, 1 of 3 PSMs at 250 fg, and 2 of 4 PSMs at 50 fg. One incorrect sequence assignment was observed at the 50 fg peptide load. Together, these results indicate that *de novo* sequencing provides a useful database-independent strategy for mutated peptide discovery, but shows less consistent sensitivity than G-PTM-D across peptides and low peptide loads.

**Figure 8.**
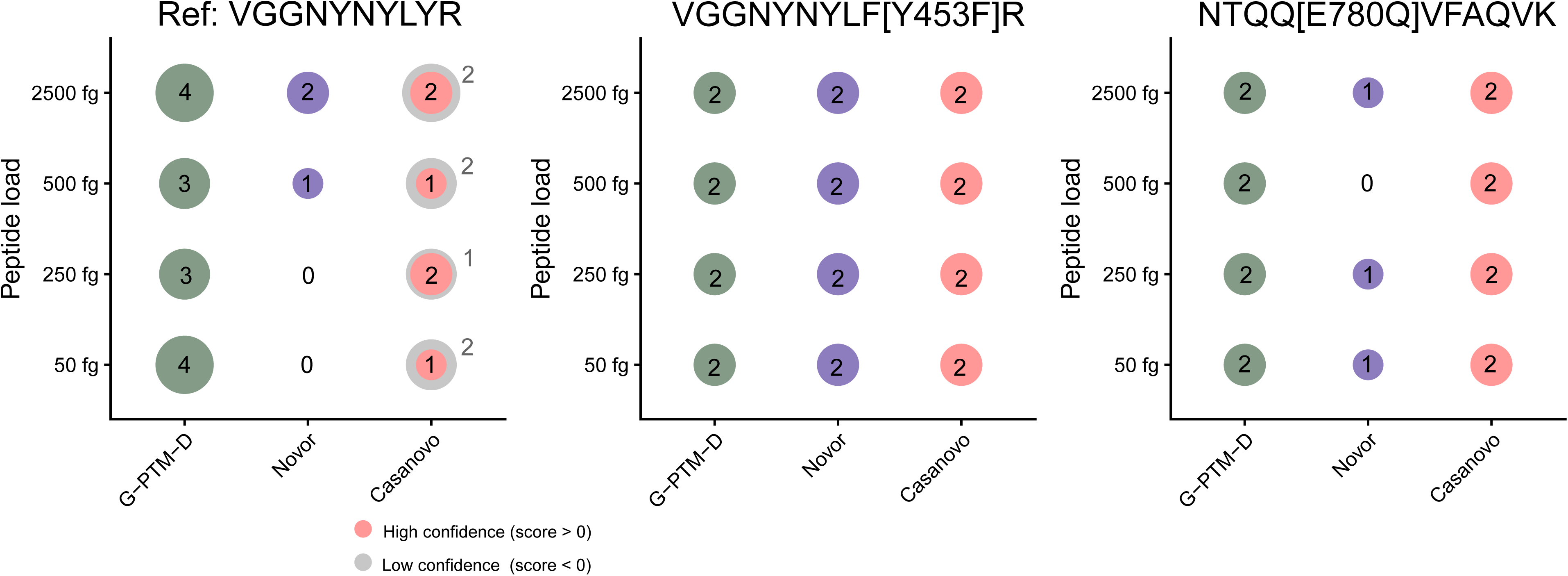
Comparison of G-PTM-D and *de novo* sequencing for peptide recovery in saliva. The bubble plot indicates the number of correctly assigned PSMs across peptide loads; absence of a bubble indicates zero correct PSMs. G-PTM-D, Novor, and Casanovo results are shown for the reference peptide VGGNYNYLYR and two single-amino-acid variant peptides, VGGNYNYLF[Y453F]R and NTQQ[E780Q]VFAQVK. Casanovo assignments are separated into higher-confidence (score > 0) and lower-confidence mass-penalized (score < 0) predictions. Novor consistently recovered VGGNYNYLF[Y453F]R but showed less consistent recovery for the other peptides, whereas Casanovo recovered the expected sequences across peptide loads with several lower-confidence assignments.

## Discussion

Advances in MS instrumentation, in particular gains in spectral acquisition rate and mass accuracy, have made deep metaproteomic profiling of complex biofluids such as saliva increasingly feasible [55, 56], creating new opportunities for simultaneous detection of viruses and their variants. Recent developments in *de novo* sequencing coupled with machine learning have demonstrated its power for peptide identification without the guidance of database in metaproteomics [54, 57, 58]. In our analysis, *de novo* sequencing performed well for mutated peptide discovery, although it remained slightly less consistent than database-correlation searching for comprehensive viral peptide detection, particularly when MS/MS spectra were affected by a noisy background, low-abundance precursor ions, or large isolation windows. Thus, for viral detection in complex saliva backgrounds, database correlation searching remains the preferred strategy because it provides greater search depth under the surveillance of a curated viral database. Nevertheless, database correlation faces fundamental statistical constraints as the search space expands in metaproteomics. Expanding the search database enlarges the candidate set considered for each spectrum, increasing the multiple-testing burden and consequently forcing more stringent score cutoffs to maintain the desired FDR. In practice, this false match penalty reduces sensitivity and can lead to missed peptide identifications, directly impacting detection of low-abundance pathogens.

Here, we developed an integrated database optimization strategy for mutation profiling of infectious viruses in saliva, focusing on RNA viruses with rapid evolution (SARS-CoV-2 and influenza H1N1/H3N2, and influenza B). The strategy combines two complementary components. First, we introduced lineage-based mutation prevalence thresholds to prioritize emerging and high-prevalence viral mutations for different viruses in a selected time window while eliminating the sequences of rare mutations that disproportionately inflate database size. Second, we applied the MetaMorpheus G-PTM-D function to discover and validate single amino-acid substitutions, thereby recovering variant peptides that are not explicitly represented in the database. Together, these elements yielded (i) a tailored mutation peptide database that mitigates sensitivity loss associated with viral mutation search, and (ii) sensitive single amino acid variants (SAAVs) detection with LODs for viral peptides.

The mutation profiling module generates a tailored mutation peptide database by directly using annotated protein sequences and applying prevalence thresholds to exclude mutations with lower expected clinical relevance. A three-month temporal window retained recently circulating viral diversity while markedly reducing database size, with 99.8% and 89.3% reductions for SARS-CoV-2 and influenza, respectively. An informative feature of the mutation prevalence data is its bimodal distribution, with maxima near 0% and 100%. This pattern is consistent with basic principles of viral evolution: most mutations arise stochastically and remain at low prevalence, while mutations that are selectively advantageous or linked to successful lineages become enriched and approach fixation. By leveraging this natural structure, prevalence thresholding removes low-prevalence mutations that are less likely to be biologically or clinically informative, resulting in substantial database reduction. A 10% prevalence threshold excluded 32,554 mutated peptides from the SARS-CoV-2 database and 101,328 mutated peptides from the combined influenza database, corresponding to reductions of 82.1% and 94.5% relative to the unfiltered databases, respectively. This reduction was especially notable for influenza viruses because of the higher mutation rate and greater sequence diversity. During database searching, the smaller database decreases PSMs assigned to decoy sequences, thereby lowering the score threshold required for 1% FDR control. This decrease in the score required to meet the 1% FDR threshold represents improved sensitivity or LODs compared with the larger database (Fig. 4E, F). Additionally, this approach is computationally efficient, straightforward to implement, and easy to interpret. A remaining concern, however, is that the viral mutation peptide databases contain substantial sequence similarity in both target and decoy entries, which may influence FDR control. To evaluate this effect, we repeated key analyses on viral lysate using shuffled decoy construction [59], which reduced sequence similarity and imposed more stringent FDR control. The negative correlation between database size and PSM identifications remained consistent. Overall PSM loss at 1% FDR increased 2.42-fold with the shuffled-decoy database compared with the reverse-decoy database, with the unfiltered 0% prevalence-threshold database showing the largest average loss of 741 PSMs, corresponding to 10.0% of control-search PSMs (Additional file 1: Fig. S5). Prevalence filtering mitigated this loss, reducing the average PSM loss to 259 PSMs at the 20% threshold and 103 PSMs at the 80% threshold, thereby retaining an additional 482 and 638 PSMs, respectively. These results indicate that prevalence-threshold-based mutation profiling helps preserve search sensitivity when stringent FDR control is applied.

The mutation discovery module utilizing G-PTM-D can recover sequence variation excluded by prevalence-based curation, albeit with some loss of sensitivity relative to direct database-guided profiling. G-PTM-D is well suited to viral mutation analysis because most GISAID-reported mutations differ by only a SAAS. Thus, G-PTM-D both restores coverage of filtered peptides and enables detection of newly emerging variants absent from the predefined database. As a quasi-open search, it infers mass shifts from the data, generates a focused candidate set, and confirms in the following database correlation while avoiding the FDR control problem that occurs when multiple organisms are searched separately from the same dataset. While it expands the search space in the initial stage, it constrains the verification stage to data-supported hypotheses, which is aligned with the goal of retaining sensitivity while controlling false positives.

The low-attomole detection of the spiked variant peptides in saliva indicates that many variant-specific peptides retain sufficient ionization and fragmentation efficiency for sensitive MS detection. These LODs are broadly consistent with previously reported targeted MS measurements of SARS-CoV-2 variant-specific peptides [31], supporting the feasible detection of variant peptides in complex matrices. Meanwhile, LODs are strongly dependent on the analytical platform, including sample preparation, chromatographic throughput, acquisition mode, and the mass spectrometer. Orbitrap Astral Zoom application-note data showed low-attomole targeted peptide sensitivity, with >92% of peptides having LODs below 25 amol and an example peptide detected at ∼6 amol on column. Both Orbitrap Astral and timsTOF Ultra studies demonstrate substantially improved peptide identification depth and metaproteomic sensitivity, with the Orbitrap Astral achieving approximately fivefold higher peptide quantification per unit time than earlier Orbitrap platforms and the timsTOF Ultra-based workflow identifying approximately fourfold more peptides than the previous timsTOF Pro workflow. [58, 60].

Although the strategy provides a comprehensive solution for known viral mutations and uncovered peptide mutation discovery, it has limitations that define key directions for improvement. At present, the mutation discovery is optimized only for single amino-acid substitutions. Peptides containing multiple mutations accounted for only a small fraction of the uncovered peptides (Fig. 5C, D). The sensitivity decreases substantially when multiple mutation sites are allowed in the G-PTM-D workflow, meaning that these peptides are not detectable for our mutation discovery module. In addition, mutations affecting protease cleavage sites pose a distinct challenge. A mutation that disrupts a cleavage site can be automatically treated as missed-cleavage peptide, whereas mutations that create new cleavage sites can generate truncated peptide sequences that are not feasible for the current database correlation algorithm. One practical mitigation is to use *de novo* sequencing to capture truncations and multi-mutation peptides, albeit typically at lower sensitivity than targeted database search. For example, the tryptic peptide VGGNYNYR, generated by the L452R substitution on SARS-CoV-2 spike protein, was successfully detected across peptide loads in the saliva matrix by both Novor and Casanovo (Additional file 2: Table S2), demonstrating the complementary value of *de novo* sequencing. A key downstream need is rigorous mutation annotation and validation. For example, sequence alignment tools are needed to identify and localize mutations. A verification method is needed to distinguish true variants from false positives, particularly when there is no evidence of the presenting species.

Variant peptide detection by MS presents well-recognized challenges for FDR control that warrant explicit discussion. Salz et al. reported that searching shotgun proteomics data from the reference cell line NA12878 with a state-of-the-art open modification search engine yielded 96.7% false-positive SAAV calls relative to long-read RNA-sequencing ground truth, even with nominal 1% target-decoy FDR control [61]. This finding highlights that, for SAAV-class peptides specifically, the FDR estimated by standard TDC may substantially underestimate the actual false discovery proportion, particularly when the search database is large, redundant, or contains many near-identical sequences. Our strategy differs from open-search variant detection in two ways that mitigate, but do not eliminate, this concern. First, our mutation peptide database is curated by prevalence and time filtering to retain peptides representing clinically prevalent viral lineages rather than every theoretically possible substitution, which reduces both database size and target-target sequence similarity. Second, the G-PTM-D module is operated as a discovery procedure followed by database-correlation verification, and not as a final classifier; mutated peptide calls are intended to support, rather than replace, orthogonal confirmation (e.g., synthetic peptide standards, viral genome sequencing, or targeted MS). Recent systematic evaluation of FDR control across MS workflows indicates that DDA database-search tools generally do control PSM-level FDR within reasonable bounds, whereas DIA tools show greater variability [62]; our use of DDA is consistent with this observation. We acknowledge that we have not performed formal entrapment-based validation of SAAV-level FDR for our viral mutation databases [62], and that variant-peptide-specific FDR validation using class-specific FDR estimation [63], or post-hoc spectral verification methods such as SpectrumAI [64] or PepQuery [65] would further strengthen confidence in individual mutation calls. In clinical or surveillance applications, variant peptide identifications produced by our strategy should be treated as candidate calls subject to orthogonal confirmation rather than as definitive single-source identifications.

Overall, our results show that mutated peptide database construction guided by the lineage-informed viral mutation analysis and data-driven amino-acid substitution discovery can maintain stringent FDR control while improving sensitivity for low-abundance variant peptides in complex saliva. This strategy provides a framework for proteomic analysis of rapidly evolving viral pathogens for clinical and research studies that is transferable to other studies in which rapid mutation is present.

## Conclusions

Our mutation profiling and discovery strategy for saliva viral proteomics balances database coverage with search sensitivity by combining prevalence-based filtering and a time window to reduce the amount of sequence added to the database to cover a rapidly mutating virus, and with single amino acid variant (SAAV) detection for emerging mutation discovery. The prevalence- and time- filtering markedly reduce database size while preserving complete coverage of high prevalence mutations. Meanwhile, SAAV detection captures the majority of low-prevalence peptides and provides a practical approach for monitoring emerging mutations using proteomics data. Collectively, this work establishes a practical and adaptable strategy for proteomics-based detection, mutation monitoring, and surveillance of rapidly evolving respiratory viruses in complex biofluid matrices.

## Methods

### Virus culture

Heat inactivated SARS-CoV-2, 2019-nCoV/USA-WA1/2020 was purchased from ATCC.

H1N1 influenza virus used in the study was generated by reverse genetics, as previously described [66]. The HA from A/Brisbane/02/2018(H1N1) IVR-190 and NA from A/Singapore/GP1908/2015(H1N1) IVR-180 were synthesized as gene fragments and cloned into the bidirectional pHW2000 reverse genetics plasmid. Virus rescue was carried out by transient transfection of HA and NA plasmids together with the plasmids for the 6 internal segments from A/Puerto Rico/8/1934(H1N1) in co-cultures of 293-T and MDCK-S cells. After rescuing, virus was propagated in 10-day-old fertilized chicken eggs and stored at -80°C. All virus stocks were sequence confirmed by Sanger sequencing [67, 68].

HSV-1 strain NS was propagated in Vero cells grown in Dulbecco’s modified Eagle’s medium supplemented with 5% fetal bovine serum. Cells were infected at a multiplicity of infection of 0.01. When the observed cytopathic effect was 100%, culture flasks were frozen at -80 °C and then thawed at room temperature. Crude virus was sonicated in an ice water bath and clarified by centrifugation at 1,400 × g for 30 mins at 4 °C. Clarified virus was pelleted by centrifugation at 72,000 × g for 90 minutes at 4 °C and resuspended in 300 µL of phosphate-buffered saline supplemented with calcium, magnesium, and HEPES (PBS++). Concentrated virus was overlayed on a 5 to 70% sucrose gradient and centrifuged at 190,000 × g for 120 minutes at 4 °C. Fractions of 0.5 mL were collected and the fraction containing peak virus titers was buffer exchanged with PBS++. The purified virus was sonicated in an ice water bath, aliquoted and frozen at -80 °C.

### Virus and synthetic peptide spike-in saliva sample preparation

Virus was spiked into pooled human saliva (Innovative Research, Inc.) using a dilution series (1×, 0.5×, 0.1×, 0.05× and 0.01×) in a final volume of 100 µL to model low-abundance clinical conditions prior to sample handling. Sodium dodecyl sulfate (SDS) was added to a final concentration of 2% (w/v). Samples were lysed by probe sonication (Branson SFX 150; probe 4C15) at 40% amplitude with 2 s on/2 s off pulses for a total of 2 min. Extracted proteins were further reduced by adding 13 µL of 100 mM dithiothreitol (DTT) and incubating at 56 °C for 40 min. Alkylation was performed by adding 4.5 µL of 550 mM iodoacetamide (IAA) and incubating in the dark at room temperature for 30 min. Excess IAA was quenched by adding 10 µL of 100 mM DTT.

Given the limited sample input, a modified solvent precipitation single-pot solid-phase–enhanced sample preparation (SP4) [69] workflow was used with carboxylate-modified magnetic beads (SpeedBead; Cytiva). Beads were washed three times with ultrapure water (Milli-Q) and collected after each wash using a magnetic rack, then resuspended to 5 µg/µL. For each sample, 4 µL of bead suspension was added and mixed thoroughly prior to addition of 300 µL ethanol. Samples were mixed on a ThermoMixer (Eppendorf) for 10 min to promote protein precipitation onto the beads. Protein–bead complexes were pelleted by centrifugation at 21,300 × g for 15 min and the supernatant was removed. Pellets were washed three times with 300 µL of 80% (v/v) ethanol; each wash was followed by centrifugation at 21,300 × g for 5 min and removal of the supernatant. After the final wash, pellets were air-dried for 2 min with tubes open.

Proteins were digested in 100 µL of 50 mM ammonium bicarbonate (ABC) containing mass spectrometry–grade Trypsin/Lys-C (Promega) at a 1:25 (w/w) enzyme-to-protein ratio, followed by overnight incubation at 37 °C. Beads were removed using a magnetic rack and the peptide-containing supernatant was collected. Peptides were dried by rotary vacuum evaporation and stored at −20 °C until UHPLC–MS/MS analysis.

For peptide spike-in experiments, lyophilized synthetic peptides (GenScript; see Supplementary Information) were resuspended in UHPLC mobile phase A (95% H₂O, 5% acetonitrile (ACN), 0.1% formic acid (FA), v/v) and quantified using the Pierce™ Quantitative Fluorometric Peptide Assay (Thermo Scientific). Synthetic peptides were spiked post-digestion into pooled saliva peptide digests generated using the workflow described above to minimize preparation-induced bias, at the desired final concentrations.

### Bulk sample preparation for virus lysate and saliva

Whole virus lysate and pooled saliva samples were processed using the same bulk preparation workflow. Viral samples corresponded to the final virus preparations described above and contained host-cell background proteins depending on the upstream propagation and handling conditions. Briefly, SDS was added to each sample to a final concentration of 2% (w/v), followed by lysis using probe sonication as described above. Protein concentrations were determined using the Pierce™ BCA Protein Assay Kit (Thermo Scientific). Reduction and alkylation were performed as described above. Proteins were then precipitated by addition of 400 µL ice-cold acetone and incubation at −20 °C overnight. Precipitated proteins were collected by centrifugation at 21,300 ×g for 15 min and washed twice with 80% (v/v) ice-cold ethanol. Protein pellets were resuspended in 100 µL of 50 mM ABC and digested as described above. Following digestion, 5 µg of peptides were desalted using C_18_ ZipTip pipette tips (Millipore) according to the manufacturer’s protocol, dried by rotary vacuum evaporation, and stored at −20 °C until UHPLC–MS/MS analysis.

### Bulk saliva peptide fractionation

Peptide fractionation was performed using in-house C_18_ solid-phase extraction (SPE) tips prepared by packing Empore™ C_18_ SPE disks into 100 µL pipette tips. Tips were conditioned with 100 µL of 80% ACN in 50 mM ABC, followed by equilibration with 50 mM ABC. 80 µg of digested saliva peptides were resuspended in 50 mM ABC and loaded onto the C_18_ SPE tip. The loaded tip was washed twice with 100 µL of 50 mM ABC to remove salts and residual contaminants. Peptides were then eluted sequentially into 16 fractions using increasing ACN concentrations in 50 mM ABC (v/v): 4%, 6%, 8%, 10%, 13%, 15%, 18%, 20%, 22%, 24%, 26%, 30%, 34%, 39%, 45%, and 70% ACN. Fractions were concatenated in a non-adjacent manner (1+9, 2+10, …, 8+16) prior to UHPLC–MS/MS.

### Ultra-high performance liquid chromatography-tandem mass spectrometry analysis (UHPLC–MS/MS)

All peptide samples were reconstituted in UHPLC mobile phase A. Peptides were separated by nano-flow UHPLC at a constant flow rate of 300 nL/min using an UltiMate 3000 nano-LC system (Thermo Fisher Scientific) with mobile phase A (95% H₂O, 5% ACN, 0.1% FA, v/v) and mobile phase B (0.1% FA in ACN).

Spike-in samples (80-min method): Peptides were eluted using the following linear gradient (%B): 1% (0–2 min), 5% (4 min), 10% (28 min), 20% (48 min), 40% (68 min), followed by a rapid increase to 90% (69–70 min). The gradient was returned to 1% B at 71 min and held at 1% B until 80 min for re-equilibration.

Bulk-study samples (140-min method): Peptides were eluted using the following linear gradient (%B): 1% (0–2 min), 5% (4 min), 25% (90 min), and 40% (120 min), followed by a rapid increase to 95% (123 min). The column was held at 95% B for 4 min, then returned to 1% B at 128 min and maintained at 1% B to the end of the run for re-equilibration.

MS data were acquired on an Orbitrap Exploris 240 mass spectrometer (Thermo Fisher Scientific) operated in data-dependent acquisition (DDA) mode. Nanoelectrospray ionization was performed using emitters from FOSSILIONTECH (The Sharp Singularity) at 2.2 kV in positive-ion mode, with the ion transfer tube temperature set to 325 °C. Full MS1 scans were acquired at 60,000 resolution over an m/z range of 400–1600 with an AGC target of 300% and a maximum injection time of 50 ms. For each MS1 scan, the top 20 most intense precursor ions were selected for MS2. Precursors were isolated using a ±1 m/z window and fragmented by higher-energy collisional dissociation (HCD) at a normalized collision energy of 30%. MS2 spectra were acquired at 30,000 resolution with an AGC target of 100%. Dynamic exclusion was set to 100 s after the first observation to reduce repeated sampling of the same precursor ions.

### In-house salivary protein database construction

Tandem MS/MS spectra were *de novo* sequenced using Novor [53] with the following settings: trypsin digestion specificity; precursor mass tolerance of 10 ppm; fragment-ion mass tolerance of 20 ppm; and the following modifications enabled during sequencing—carbamidomethylation (C), deamidation (N), and oxidation (M). *De novo* peptide candidates were mapped to protein sequences in Swiss-Prot to obtain protein accessions. The resulting accessions were merged with protein lists tailored from prior salivary metaproteomics studies [2–4], relevant clinical-condition saliva proteomics studies [70–72], and human salivary protein database [73]. Protein sequences for the combined accession set were retrieved from Swiss-Prot and compiled into a FASTA database using a custom R script, yielding 14,017 protein sequences.

### Peptide database and decoy construction and representativeness validation

SARS-CoV-2 viral population between June 1 and August 31, 2025 were downloaded with annotated mutation sites from EpiCoV^TM^ at GISAID. Mutated protein sequences were generated by applying the reported amino-acid substitutions to the Wuhan reference genome (NC_045512.2) using a custom R script. For influenza A (H1N1, H3N2) and influenza B, virus entries between November 1, 2024 and February 28, 2025 were downloaded from EpiFlu^TM^ at GISAID as protein sequences. All viral protein sequences were digested *in silico* using trypsin specificity with up to two missed cleavages. Candidate peptides were filtered to 5–30 amino acids to match typical bottom-up proteomics observability. Peptides were stratified by lineage for SARS-CoV-2 and by clade for influenza. For each lineage (SARS-CoV-2) or clade (influenza), peptide prevalence was computed as:

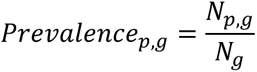

Equation 1. Peptide prevalence within a lineage/clade. For peptide 𝑝 in lineage/clade 𝑔, 𝑁_𝑝,𝑔_ is the number of viral sequences in 𝑔 that contain peptide 𝑝, and 𝑁_𝑔_ is the total number of viral sequences assigned to lineage/clade 𝑔.

A prevalence threshold was then applied within each lineage/clade to remove low-prevalence peptides. To avoid redundancy across groups, peptides shared between multiple lineages/clades were deduplicated (i.e., identical peptide sequences were retained once). The resulting non-redundant peptide set was exported as the target (mutated) peptide FASTA database.

### Peptide decoy database construction

A decoy database was generated from the target peptide set using two decoy-generation approaches. For the primary decoy set, reversal decoys were created by retaining the C-terminal tryptic residue (K/R) and reversing the remaining residues; peptides that did not end in K/R were reversed in full. For a more stringent decoy set, internal-residue shuffling was performed by keeping the N- and C-terminal residues fixed and randomly shuffling the internal residues. For both approaches, decoy peptides were required to be unique, and the final decoy database was constructed to be the same size as the target database.

### Validation of database representativeness at different thresholds

Database representativeness was validated by calculating the coverage of upcoming viral entries. An independent “next-month” viral population was retrieved from GISAID (September 1–30, 2025 for SARS-CoV-2; March 1–31, 2025 for influenza A H1N1/H3N2 and influenza B). These sequences were processed using the same *in silico* digestion and peptide length filtering workflow described above. Each resulting tryptic peptide was then labeled as covered if its sequence was present in the mutated peptide database and uncovered otherwise. Uncovered peptides were further classified into three categories: (i) single amino-acid substitutions that did not involve K/R, (ii) single amino-acid substitutions involving K/R, and (iii) all other remaining cases. Coverage was quantified for peptide databases generated under different prevalence thresholds, and analyses were performed both across the full viral population and within specific viral proteins.

### MS data processing and peptide identification

Conventional database searching was performed in MetaMorpheus (v1.0.8) using a target–decoy strategy with results filtered at 1% FDR. Searches assumed trypsin specificity. The precursor mass tolerance was set to 10 ppm and the product-ion mass tolerance to 20 ppm. Carbamidomethylation on C was specified as a fixed modification and oxidation on M as a variable modification. All other parameters were left at MetaMorpheus defaults unless otherwise noted. Database searching was also performed in FragPipe using settings comparable to those applied in MetaMorpheus, followed by peptide-spectrum match validation with PeptideProphet.

G-PTM-D searches were performed in MetaMorpheus with a precursor mass tolerance of 10 ppm and a product-ion mass tolerance of 20 ppm. The maximum number of modifications per peptide prior to the G-PTM-D step was set to 2. Carbamidomethylation on C was specified as a fixed modification and oxidation on M as a variable modification. The G-PTM-D modification list included all documented amino-acid substitutions consistent with single-nucleotide and multi-nucleotide changes (Additional file 2: Table S3),.

To perform a regular PTM-style search that enumerates possible single amino-acid substitutions, the search parameters were identical to the conventional database search described above, except that the variable modification set was expanded to include all documented substitution masses as mentioned above.

*De novo* sequencing by Novor and Casanovo of spike-in saliva samples used the same settings as applied for *de novo* sequencing of the saliva fractionation experiments, with carbamidomethylation on C specified as a fixed modification and oxidation on M as a variable modification to match the G-PTM-D task settings.

## Declarations

### Ethics approval and consent to participate

Not applicable

### Consent for publication

Not applicable

### Data availability

The MS proteomics data have been deposited to the ProteomeXchange Consortium via the PRIDE [74] partner repository with the dataset identifier PXD077745.

### Authors’ contributions

Y.Z. designed the workflow, prepared samples, constructed databases, analyzed LC-MS/MS data, generated figures, and drafted the manuscript. M.S. contributed to data analysis, and manuscript editing. A.T. supervised the study and revised the manuscript. All authors read and approved the final manuscript.

## Acknowledgements

We gratefully acknowledge Dr. Jefferson Santos and Prof. Scott E. Hensley for providing the H1N1 samples, and Dr. Kevin Egan and Prof. Harvey M. Friedman for providing the HSV-1 samples used in this study.

## Supplementary information

### Additional file 1

Figure S1: Prevalence distribution of SARS-CoV-2 mutations from viral population sequences collected between June 1 and August 31, 2025.

Figure S2: A, B, PSM identification loss from cultured SARS-CoV-2 (A) and influenza A H1N1 (B) samples compared to the viral reference sequences and contaminants by FragPipe.

Figure S3: A, B, PSM identification loss from virus spiked into saliva: SARS-CoV-2 (A) and influenza A H1N1 (B) samples compared to the viral reference sequences and contaminants by MetaMorpheus.

Figure S4: A, Heat map of HSV-1 peptides detected in a saliva spike-in sample with 1× dilution factor, no detection in 0.5×, 0.1×, 0.05×, and 0.01x samples. B, Log-log calibration curve for the reference peptide VSGGPGPLVLR in saliva matrix (R² = 0.9624). C, Mass fractions of HSV-1 proteins determined by the total protein approach.

Figure S5: PSM identification loss from SARS-CoV-2 viral lysate using shuffled decoy database with different prevalence thresholds (0%, 20%, 80%).

### Additional file 2

Table S1. Comparison of PSM counts from final database correlation using a G-PTM-D-generated mutation database and a customized mutation-inclusive database.

Table S2. SARS-CoV-2 spike protein mutated peptides VGGNYNYR [L452R] detection by Novor and Casanovo. (n=2)

Table S3. Single amino acid substitutions for G-PTM-D and conventional PTM search

