## Supplementary figures and images for "A Viral Mutation Profiling and Discovery Strategy for Sensitive Multiplex Detection of Viruses and Variants in Saliva by Proteomics"

### Additional File 1

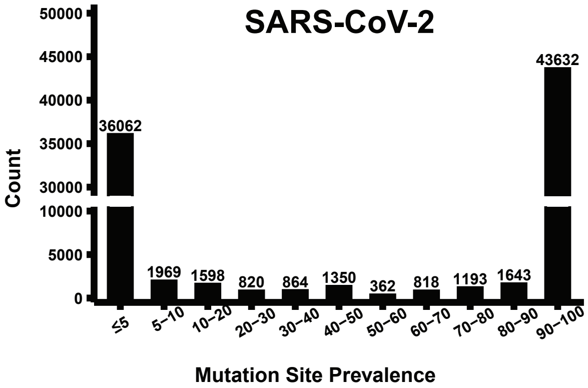

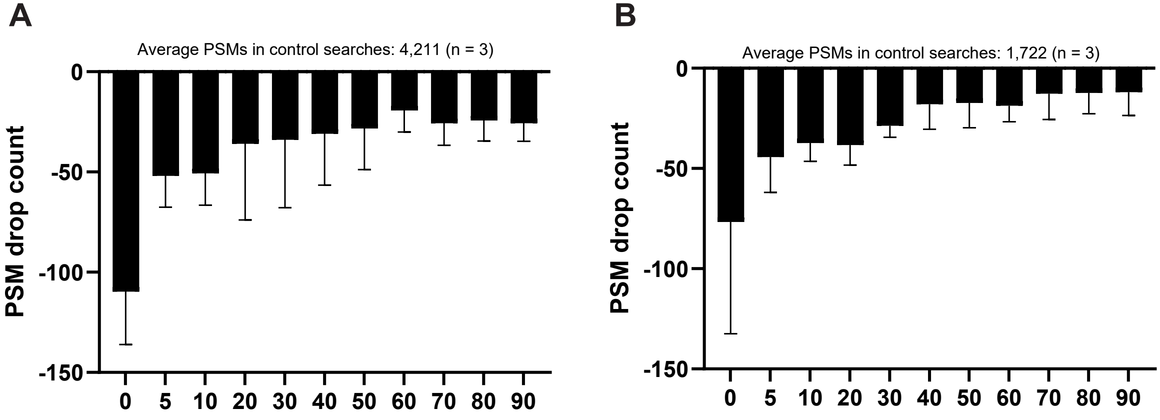

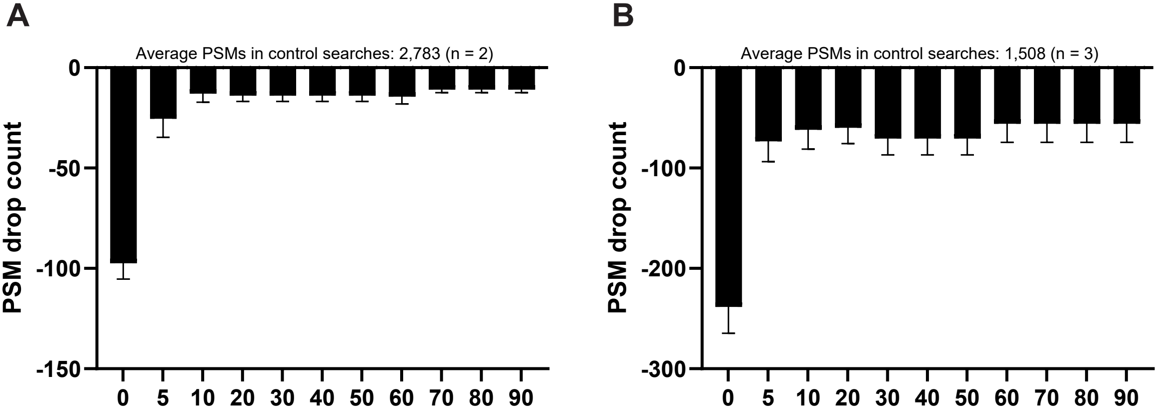

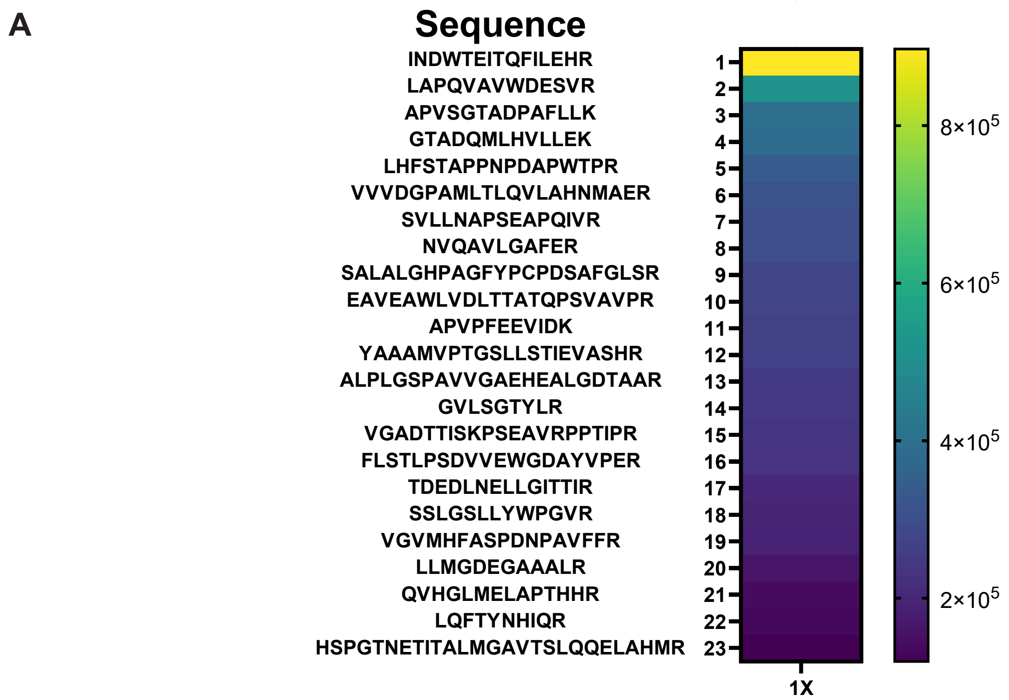

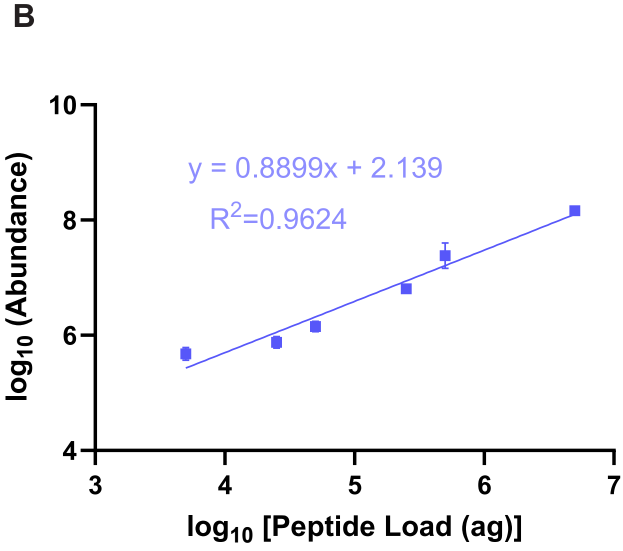

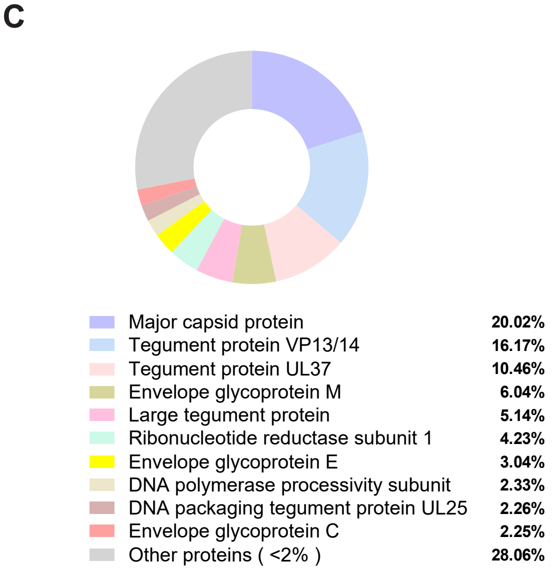

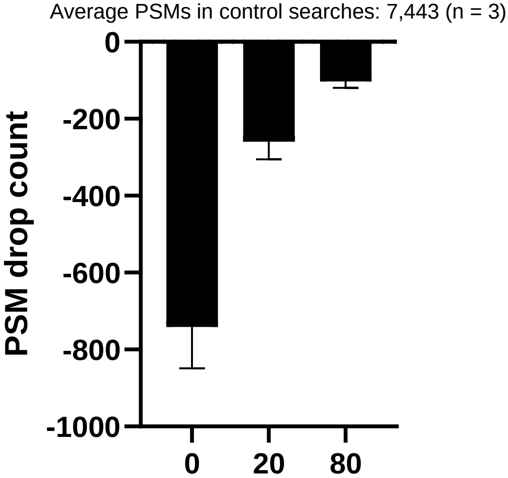
